## Supplementary figures for "Interrogation of RNA-protein interaction dynamics in bacterial growth"

#### Affiliations

#### Author List footnotes

* Lead contacts

#### Correspondence

Anne E. Willis (aew80 @ mrc-tox.cam.ac.uk), Kathryn S. Lilley (k.s.lilley @ bioc.cam.ac.uk) & Eneko Villanueva (ev318 @ cam.ac.uk)

### Extended Data Figures

#### Extended Data figure 1

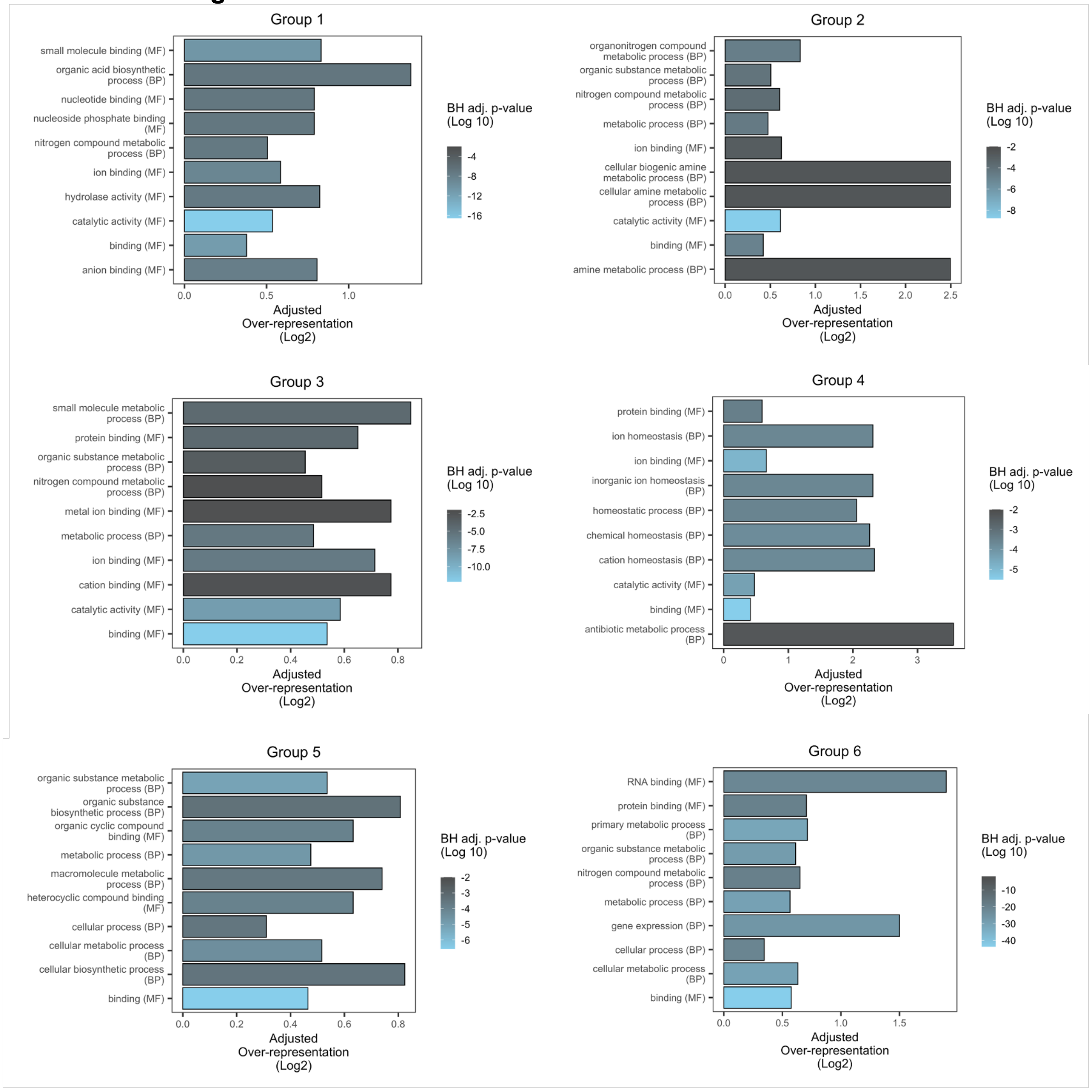


**Figure S1: GO-term over-representation at specific growth stages.** Top ten molecular function (MF) or biological process (BP) GO terms over-represented in proteins identified in each cluster across the growth curve. BH adj. P-value: Benjamini-Hochberg adjusted P-value.

#### Extended Data figure 2

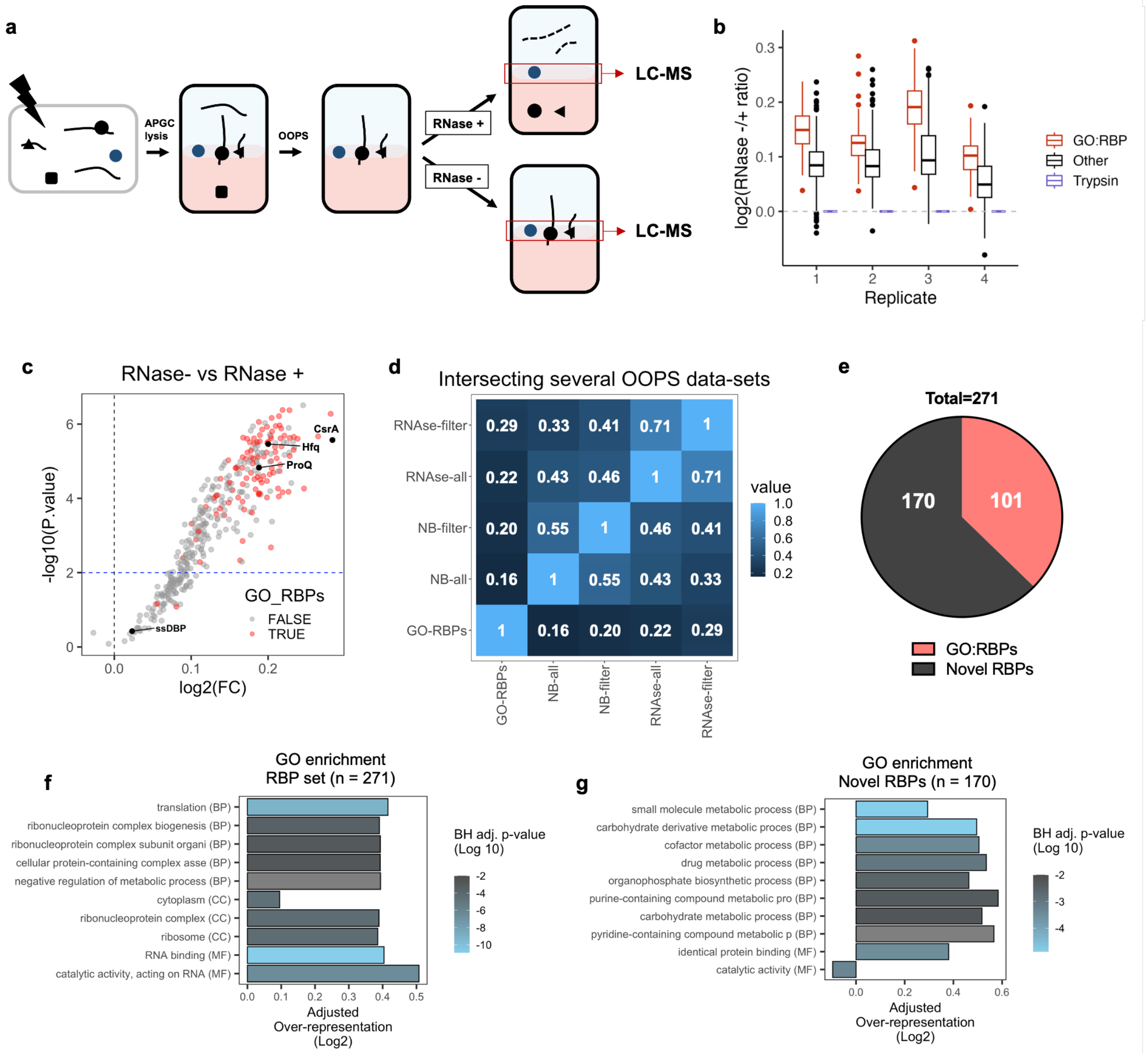


**Figure S2: RNase assay to determine *E. coli* RBPome.**
**a**, Schematic representation of the OOPS method to catalogue RBPs. APGC: Acid-phenol guanidium chloroform. RNA depicted as strings and proteins as solid colours. **b**, MS quantitation of relative protein abundance between RNase negative and RNase positive samples. **c**, Volcano plot of RNase-/+ ratios. CsrA, Hfq and ProQ are highlighted as representative RBPs and single-stranded DNA-binding protein (ssDBP) is a negative control. GO annotated RBPs are highlighted in red. **d**, Proportion of intersection with other RBP sets including GO-RBPs (180 proteins), proteins annotated as RNA-binding (‘GO:0003729’); NB-all (655), all proteins identified in bacterial RBPome from Queiroz et al.; NB-filter (364), filtered bacterial RBPome set from Queiroz et al.; RNase-all (382), all proteins identified in RNase assay; RNase-filter (271 proteins), proteins with adjusted P-value < 0.01 in RNase assay. **e**, Number of RBPs with annotated RNA-binding function. **f**, GO-term over-representation analysis of all 271 RBPs. **g**, GO-term over-representation analysis of novel RBPs (170 proteins). GO term enrichment conducted against all proteins identified in MS experiment.

**Extended Data figure 3**

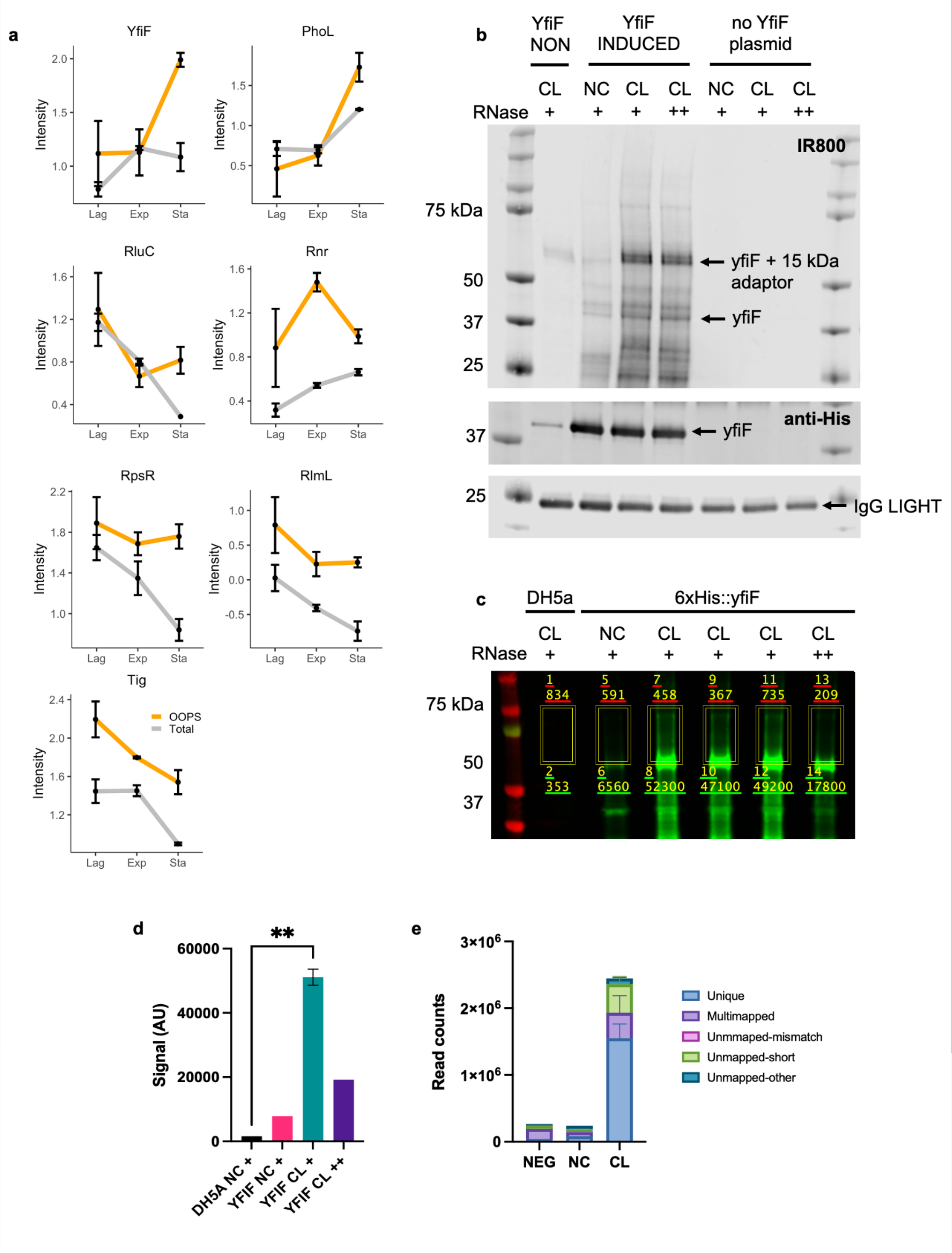


**Figure S3: Validation of YfiF as novel RNA-binding protein. a,** Protein abundance levels for YfiF and physical interactors, data shown as mean +/- SD across samples. In orange, OOPS extraction type; grey, total abundance extraction. **b**, PNK assay for YfiF. '+', refers to 1:300 RNase dilution, '++', refers to 1:100, 'YfiF NON', non-induced YfiF-His-tag transformed strain; 'YfiF INDUCED', induced YfiF-His-tag transformed strain; no YfiF plasmid, untransformed DH5α strain. RNA adaptor for PNK assay has a molecular weight of 15 kDa.

**c**, Left: Repeat PNK assay for YfiF target, '+' refers to 1:6000 RNase dilution, '++' refers to 1:100. **d**, Densitometry quantification of IR800 fluorescent adaptor, area quantified shown on gel in **c**. These areas indicate regions where CL RNA was extracted for iCLIP. Significantly more signal seen between the CL and NC (negative control) samples (*P*-value: 0.0034, Student's t-test). **e**, Absolute reads counts of each sample according to alignment of the read by STAR.

**Extended Data figure 4**

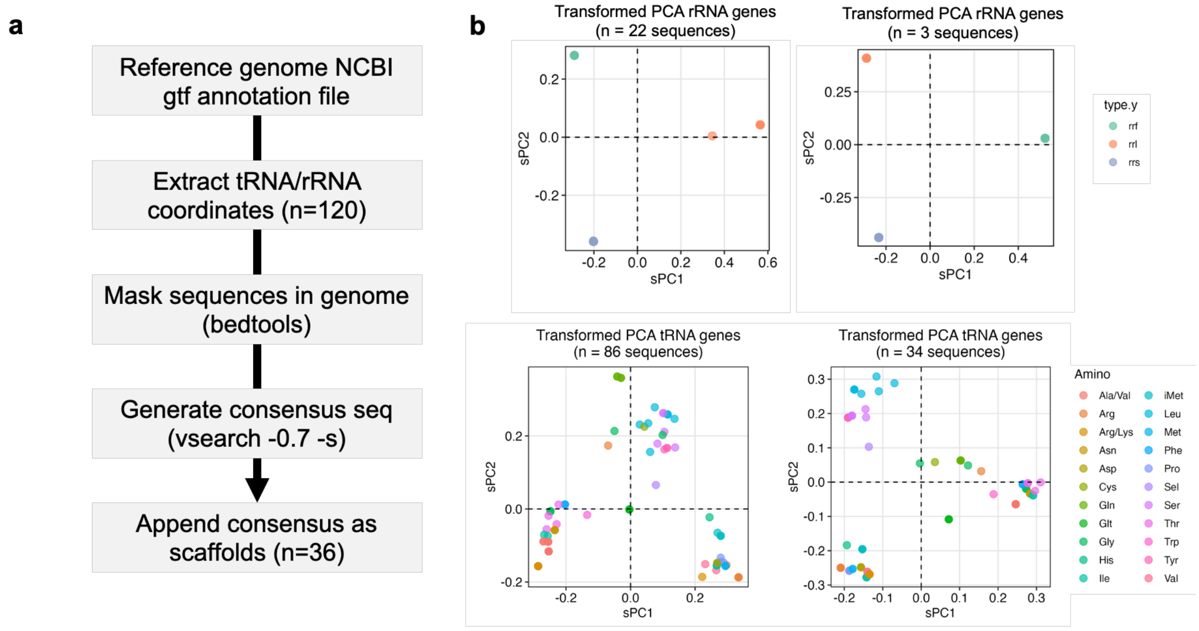


**Figure S4: Generation of a non-redundant genome for iCLIP data analysis. a,** Schematic representation of *E. coli* artificial genome for alignment. The coordinates for tRNA and rRNA genes were extracted from the gtf annotation file and masked in the genome. The extracted fasta sequences were clusterized to 70% similarity to generate consensus sequences which were then appended as scaffolds to the masked genome. **b,** Left: PCA plots of individual rRNA/tRNA genes in the *E. coli* genome. Right: PCA plots of consensus rRNA/tRNA genes.

#### Extended Data 5

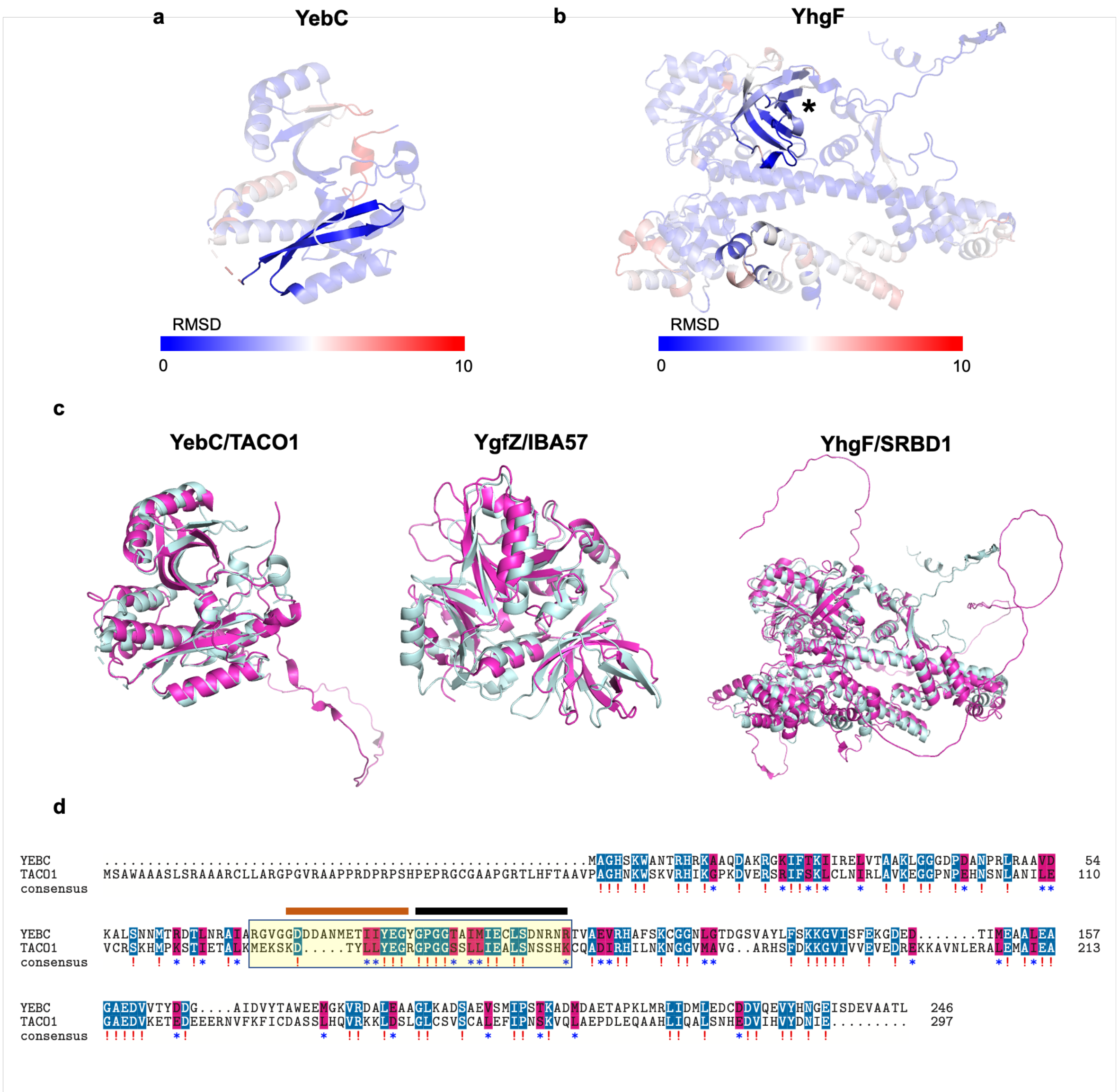


**Figure S5: Structural conservation between experimentally unannotated proteins and human orthologs. a,** YebC (1KON) and its human ortholog (TACO1; AF-Q9BSH4-F1) aligned and coloured by RMSD. Only YebC shown here. Highlighted in opaque is the predicted RNA-binding site by RBDpep[23] and/or OOPS[4]. **b**, YhgF (AF-P46837-F1) and its human ortholog (SRBD1, AF-Q8N5C6-F1) aligned and coloured by RMSD. Dark blue is good alignment, higher deviations are in red. Residues not used for alignment are coloured grey. Highlighted in opaque is both the S1 RNA binding domain (signalled with an asterisk) and the predicted RNA-binding site by RBDpep[23] and/or OOPS[4]. **c**, Full protein structure alignments between *E. coli* (cyan) and *H. sapiens* (fuchsia). YebC/TACO1: 1KON/AF-Q9BSH4-F1; YgfZ/IBA57: 1VLY/6QE3; and YhgF/SRBD1: AF-P46837-F1/AF-Q8N5C6-F1. **d**, Pairwise alignment between YebC and TACO1 amino acid sequences. Boxed in yellow is the LysC peptide identified by both OOPS[4] and RBDpep[23] as involved in RNA interaction. In orange, this specific region is predicted to bind RNA by OOPS, in black by RBDpep.

#### Extended Data Figure 6

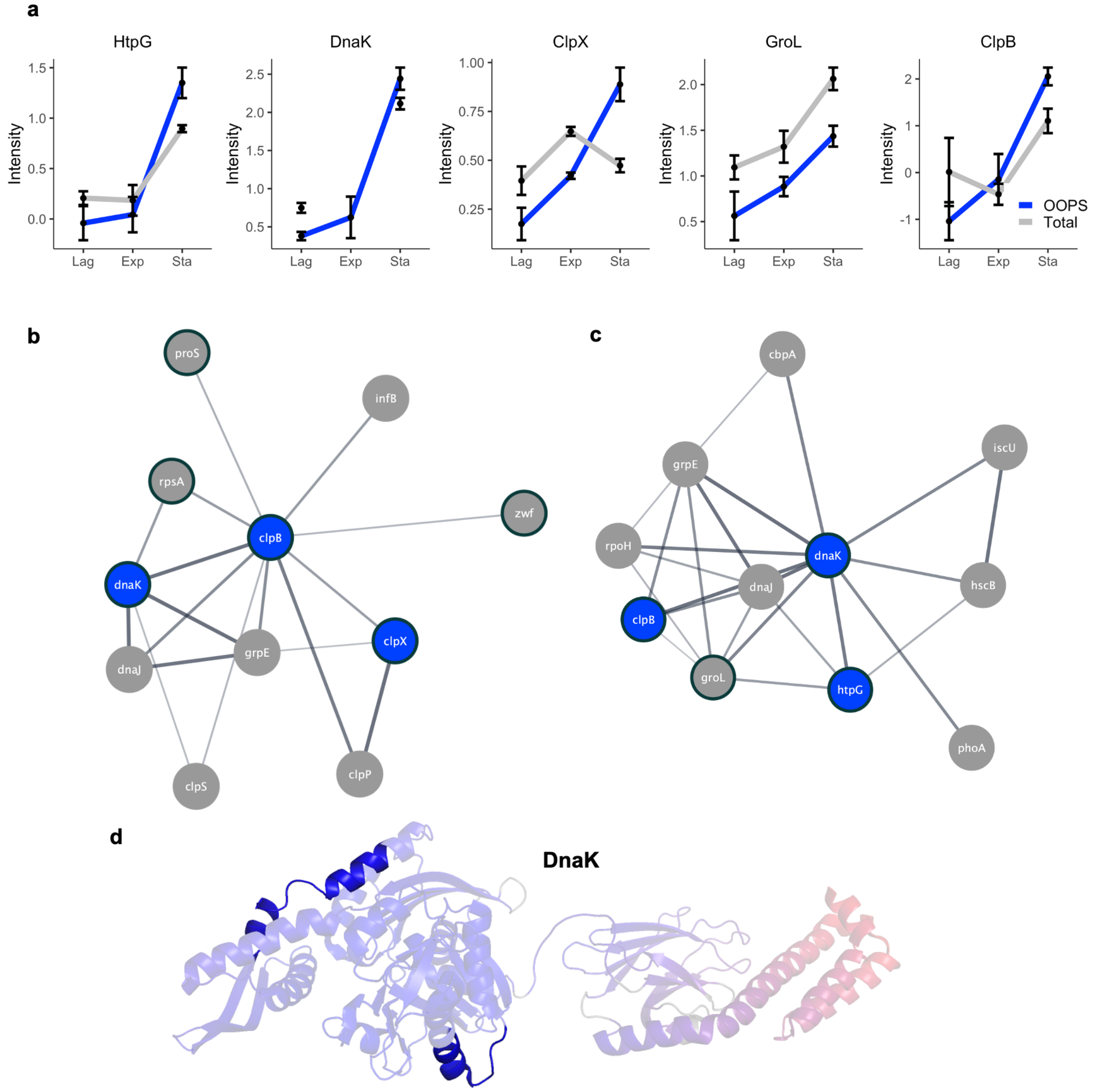


**Figure S6: Protein chaperones display consistent RNA-binding profiles.**
**a,** Protein abundance levels for protein chaperones identified with increased RNA-binding profile in the stationary phase, data shown as mean +/- SD across samples. In blue, OOPS extraction type; grey, total abundance extraction. **b-c**, Physical interaction network of ClpB (b) and DnaK (c) as annotated in STRING-db. In blue, proteins that significantly bind more RNA in the stationary phase. Bold outline highlights proteins identified as RBP via the RNase assay. **d**, NMR structure of DnaK protein (PDB 1D: 2KHO) coloured by RMSD score when aligned with the human ortholog (HSPA9). In opaque are the peptides predicted to interact with RNA in human orthologs by[4,32].

**Extended Data Figure 7**

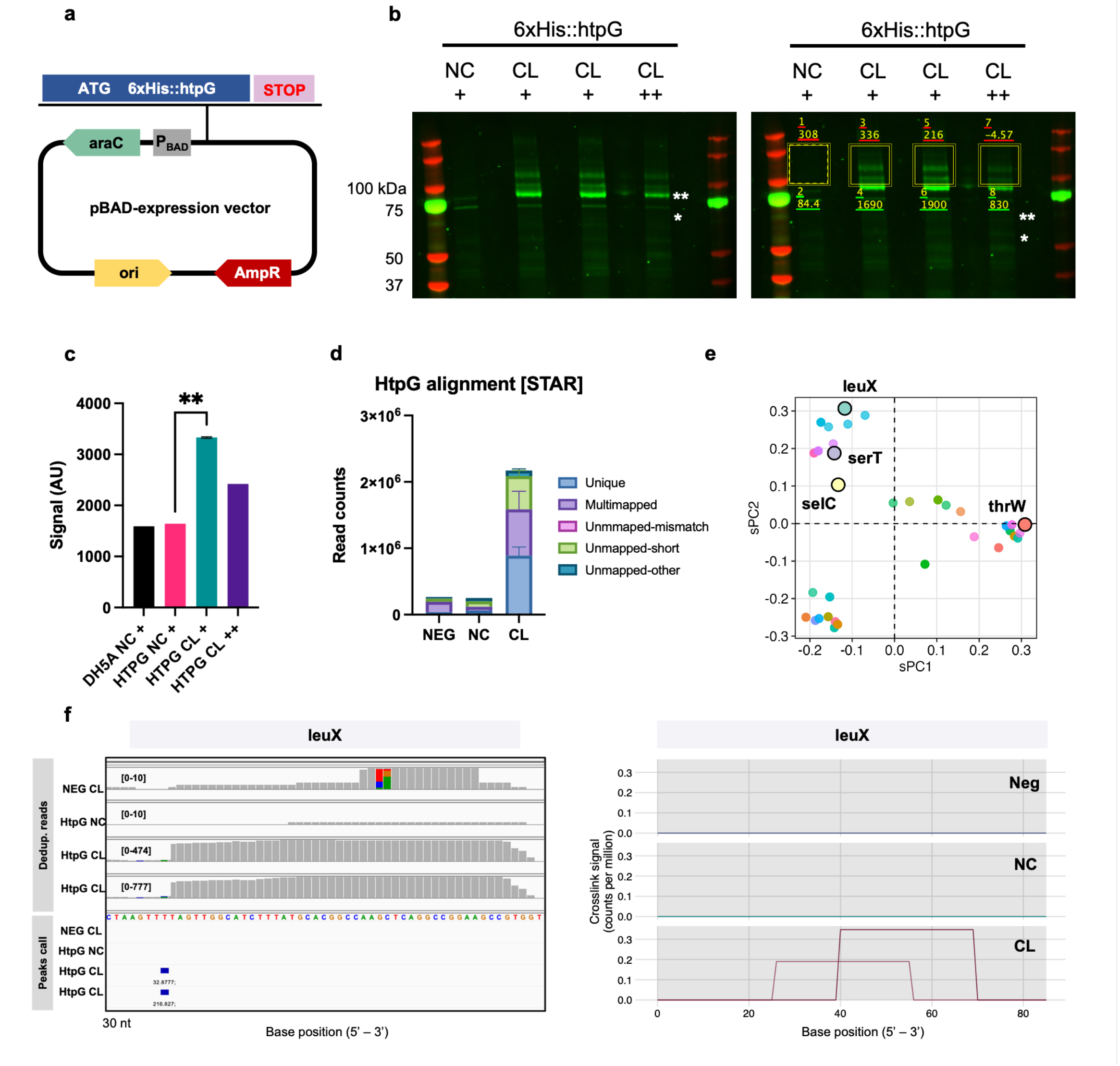


**Figure 7: Protein chaperones as RNA-binding proteins.**

**a,** Schematic representation of transformed plasmid with N-terminus tagged htpG under an arabinose inducible promoter. **b,** PNK assay with three technical replicates. On the right, boxed regions indicate the area of gel from which CL RNA was extracted for iCLIP per sample. One asterisk (*) indicates predicted molecular weight of HtpG, two asterisks (**) indicate predicted molecular weight of HtpG + 15 kDa adaptor. NC: non-crosslinked, CL: crosslinked, '+': RNase digestion 1:6000, '++': RNase digestion 1:100. **c**, Densitometry quantification of IR800 fluorescent adaptor, area quantified shown on gel. Significantly more signal seen between the CL and NC (negative control) samples (*P*-value: 0.0065, Student's t-test). **d**, Absolute reads counts of each sample according to alignment of the read by STAR. **e**, Location of HtpG tRNA targets on sequence-based PCA. **f**, Top: de-duplicated reads against leuX gene in IGViewer. Bottom: Crosslink sites ('peaks') identified by PureCLIP and their associated score. **g**, Analysis of iCLIP datasets mapped to leuX targets by iCount in agreement to e. Crosslink counts are visualised and normalised to library size with CLIPplotR.
